# Integrin Activation Enhances Lesion-Specific Targeting of Monocyte-Mimetic Nanoparticles in Atherosclerosis

**DOI:** 10.64898/2026.03.04.707824

**Authors:** Ting-Yun Wang, Jiapei Jiang, Joshua Rousseau, Zijian Wan, Kylie Hartana, Shaopeng Wang, Kuei-Chun Wang

**Author notes:** Corresponding author: Kuei-Chun Wang, Ph.D., 501 E. Tyler Mall, Engineering Center G Wing, 346 Tempe, AZ 85287-9709.

## Abstract

**Purpose:** Endothelial cell (EC) activation, characterized by upregulation of adhesion molecules that drive monocyte recruitment, contributes to plaque progression while also providing an opportunity for targeted therapeutic delivery. Leveraging the cell membrane cloaking strategy, we recently developed a monocyte-mimetic nanoparticle (MoNP) platform that exploits the natural inflammatory tropism of monocytes for site-specific delivery to atherosclerotic vessels. Recognizing that integrin activation is a key determinant of monocyte adhesion to ECs, this study investigates whether pre-activating integrins on MoNP enhances their binding affinity and accumulation at atherosclerotic lesions.

**Methods:** Mouse bone marrow-derived monocytes were pretreated with CCL2 or Mn^2^□ to activate membrane integrins. Isolated monocyte plasma membranes were cloaked onto fluorescently labeled polymeric cores to generate integrin-activated MoNPs (IA@MoNPs). The targeting capability of IA@MoNPs toward endothelial ligands, inflamed ECs, and atherosclerotic lesions was evaluated using *in vitro* and *in vivo* models.

**Results:** IA@MoNPs exhibited markedly enhanced binding to VCAM1, the primary endothelial ligand mediating integrin-dependent monocyte adhesion, and significantly increased uptake by ECs under atheroprone conditions compared to standard MoNPs. *In vivo*, IA@MoNPs demonstrated enhanced accumulation in atherosclerotic arteries without increasing nonspecific binding, and blocking β1-integrins on IA@MoNPs abolished this targeting effect. Importantly, integrin activation on IA@MoNPs did not compromise circulatory stability or induce immune or organ toxicity.

**Conclusion:** Integrin activation represents a simple yet effective strategy to enhance MoNP targeting to inflamed ECs and atherosclerotic lesions. This mechanism-driven approach improves targeting performance while maintaining specificity and safety, advancing the translational potential of the biomimetic nanomedicine platform for atherosclerosis.

## 1. Introduction

Atherosclerosis, driven by chronic inflammation within the arterial wall, remains the primary cause of life-threatening cardiovascular events [1, 2]. Endothelial cells (ECs), which line the innermost layer of blood vessels, typically reside in a quiescent state, maintaining a non-adhesive interface with circulating blood cells [3, 4]. However, exposure to atheroprone stimuli, including pro-inflammatory chemokines and cytokines, oxidized low-density lipoprotein (oxLDL), and disturbed flow, triggers EC activation, characterized by increased expression of adhesion molecules that mediate monocyte recruitment and subsequent macrophage infiltration into the vessel wall [4]. The sustained accumulation of macrophages and their lipid uptake lead to foam cell formation, driving the development and progression of atherosclerotic plaques. While EC activation is central to this inflammatory cascade, the disease-specific molecular signatures displayed by inflamed ECs may provide an avenue for targeted therapeutic delivery to atherosclerotic lesions. Such targeted strategies could potentially modulate pathological events within plaques and complement existing treatments that primarily focus on managing cardiometabolic risk factors.

Nanoparticles have gained significant attention as a strategy for site-specific drug delivery [5]. Selective targeting is often achieved by conjugating nanoparticles to ligands, such as peptides, antibodies, and glycans, that bind disease-associated markers [6, 7]. In atherosclerosis, growing efforts have focused on nanoparticles functionalized with ligands that target adhesion molecules overexpressed by inflamed ECs, including vascular cell adhesion molecule 1 (VCAM1) and intercellular adhesion molecule 1 (ICAM1) [8–11], as well as extracellular matrix components such as fibronectin [12]. However, significant challenges, including potential immunogenicity, short circulation time, and the requirement for high-avidity ligands or multivalent interactions to achieve robust targeting remain barriers to clinical translation [13].

Cell membrane cloaking has emerged as a promising strategy to functionalize nanoparticles, offering biomimetic properties inherited from source cells that improve circulation stability and enable site-specific delivery [14, 15]. Leveraging this approach, we established a monocyte mimetic-nanoparticle (MoNP) platform that harnesses the monocyte membrane-derived receptors to directly bind adhesion ligands on inflamed ECs lining atherosclerotic plaques [16]. We have demonstrated enhanced binding to and uptake of MoNPs by cytokine-activated ECs *in vitro*, and preferential accumulation in atherosclerotic arteries *in vivo*. Furthermore, MoNPs can safely and effectively deliver therapeutic and imaging agents, leading to enhanced treatment efficacy and improved detection sensitivity for plaques [16, 17]. Despite these promising findings, prior MoNPs were prepared with membranes from naïve monocytes that do not fully capture the adhesive phenotype of activated monocytes. To enhance biomimicry and lesion-specific targeting, we re-engineer MoNPs by incorporating monocyte membranes enriched in activated targeting moieties to maximize binding affinity and specificity to atherosclerotic vessels.

Integrins expressed on monocyte membrane, particularly very late antigen-4 (VLA-4; integrin α4β1) and macrophage-1 antigen (Mac-1; αMβ2), play a central role in mediating monocyte-endothelial interactions [18–20]. Upon activation by pro-inflammatory cues that induce conformational changes, these receptor integrins bind their respective endothelial ligands, VCAM1 for VLA-4 and ICAM1 for Mac-1, facilitating firm monocyte adhesion on inflamed ECs. To determine whether these mechanisms can be leveraged to enhance plaque targeting, we investigate two strategies to activate integrins on monocyte membranes prior to MoNP formulation: pretreatment of monocytes with C-C motif chemokine 2 (CCL2) and exposure to manganese ions (Mn^2+^). CCL2, a chemokine abundantly present in atherosclerotic environments, binds to its receptor CCR2 on monocytes to trigger inside-out signaling that increases integrin activation, clustering, and affinity for endothelial ligands [21, 22]. In parallel, Mn^2+^ is a well-established pan-integrin activator that directly engages in metal ion-dependent adhesion sites within integrins, inducing and stabilizing their high-affinity conformations [23, 24]. These integrin activation approaches provide a mechanistic basis for re-engineering MoNPs with greater adhesive capacity, enabling more effective targeting of atherosclerotic plaques.

To implement the integrin activation strategy, we pretreat mouse monocytes with CCL2 or Mn^2+^ to activate membrane integrins prior to cloaking them onto fluorescently labeled polymeric cores, generating integrin-activated MoNPs (IA@MoNPs). We first evaluate IA@MoNPs binding to VCAM1 and to ECs exposed to atheroprone stimuli *in vitro*, assessing their binding affinity and uptake in comparison with standard MoNPs formulated with naïve monocyte membranes. We then assess targeting performance *in vivo* using a murine model of carotid atherosclerosis. Additionally, we examine how pre-activating membrane integrins on MoNPs influences protein corona formation, circulation stability, and safety profiles. Collectively, this study establishes a simple, mechanistically informed strategy to further augment MoNP-mediated delivery to inflamed endothelium, advancing the translational potential of biomimetic nanocarriers for atherosclerotic disease.

## 2. Materials and Methods

### 2.1. Cell culture and monocyte membrane isolation

Primary monocytes used in this study were derived from bone marrow cells (BMCs) isolated from C57BL/6 mice as previously described [16]. BMCs were flushed from the tibiae and femurs, then cultured in RPMI 1640 medium (Cytiva #SH30027) supplemented with 10% FBS, 1% penicillin-streptomycin, 1% L-glutamate, 1% sodium pyruvate, and 20 ng/mL M-CSF (Biolegend #576406) for 5 days to induce differentiation into monocytes. Monocytes were subsequently sorted using the MojoSort™ Mouse Monocyte Isolation Kit (BioLegend #480154), followed by plasma membrane isolation using the cell fractionation kit (Invent Biotechnologies). To activate integrins, monocytes were pre-treated with 100 ng/mL of CCL2 (Biolegend #578402) or 3 μM of MnCl_2_ for 30 minutes prior to membrane isolation.

Human umbilical vein ECs (HUVECs), purchased from ATCC (#PCS-100-013) and maintained in the Endothelial Cell Grown Medium (Cell Application #211-500), were used for *in-vitro* nanoparticle testing. All cells were maintained under standard cell culture conditions (37ºC, 5% CO_2_, 100% humidity).

### 2.2. Nanoparticle formulation

MoNPs were prepared as previously described [16]. Briefly, poly(D, L-lactide-co-glycolide) (PLGA, Sigma Aldrich #Resomer® RG 503H), with or without a fluorescent dye DiD (Biotium), were dissolved in dichloromethane (Sigma-Aldrich). The organic phase was added dropwise to a poly(vinyl alcohol) (PVA, Acros Organics) solution under probe sonication. The resulting emulsion was transferred into additional PVA solution to allow solvent evaporation. After repeated washing and centrifugation, PLGA nanoparticle cores (bare NPs) were collected and subsequently cloaked with either integrin-activated or naïve monocyte membranes to generate IA@MoNPs or MoNPs, respectively, by mixing and sonicating the membrane-NP suspension in an ice-water bath. Their physicochemical properties, including hydrodynamic diameter and zeta-potential, were measured using dynamic light scattering (DLS, Malvern Panalytical). Membrane proteins were assessed by Western blot using antibodies against CD11b (Cell Signaling #17800, 1:1000), Na^+^/K^+^ ATPase (Cell Signaling #3010, 1:1000), α4-integrin (Invitrogen #PA5-20599, 1:1000), and β1-integrin (Cell Signaling #4706S, 1:1000).

### 2.3. Monocyte adhesion strength assay

Monocytes labeled with CellBrite® Dye (Biotium # 30021) were stimulated with CCL2, Mn^2+^, or PBS and then incubated with ECs to assess their binding affinity. Fluorescently labeled monocytes were added to a monolayer of TNFα-activated ECs seeded on a collagen I-coated slide and allowed to attach for 30 minutes under static conditions. After washing off unbound cells, initial fields of view (FOVs) were recorded using the Lionheart FX Automated Microscope. The slides were then mounted in a parallel plate flow chamber [25], and ECs were subjected to increasing laminar shear stresses (SS): 1 dyne/cm^2^ for 2 minutes, followed by imaging the same FOVs, subsequently 6 and 12 dyne/cm^2^ using the same procedure. Bound monocytes remaining after shear exposure were quantified using ImageJ and normalized to the initial binding observed under static conditions.

### 2.4. VLA-4 activation assays

VLA-4 activation on CCL2, Mn^2+^, or PBS stimulated monocytes were evaluated using the LDV-FITC probe (Tocris Bioscience #4577), which binds to the activated conformation of VLA-4. CCL2-, Mn^2+^-, or PBS-stimulated monocytes were incubated with LDV-FITC, and unbound LDV-FITC was removed by centrifugation. These monocytes were then resuspended in ultrapure water, and fluorescence intensity was quantified using the Attune NxT Flow Cytometer (ThermoFisher) at the ASU Flow Cytometry Core Facility. Enhanced LDV-FITC fluorescence was interpreted as enhanced VLA-4 activation.

To access functional binding, 100 μg of IA@MoNPs, MoNPs, or bare NPs were incubated with 2 μg of recombinant VCAM1 (PeproTech #315-37). After washing, VCAM1-bound to nanoparticles were analyzed by Western blot using antibodies against VCAM1 (Cell Signaling #13622, 1:1000) and CD11b (Cell Signaling #17800, 1:1000). Protein bands were detected using a chemiluminescence kit (Pierce) and imaged with an Analytik Jena Gel-Doc system and quantified using ImageJ to compare VCAM1 binding between nanoparticles.

Binding strength to VCAM1 was further evaluated using a modified surface plasmon resonance (SPR) assay [26, 27]. A carboxylated gold surface was activated using NHS/EDC for 15 minutes at 5□μL/min, followed by immobilization of recombinant VCAM1 (1□μM) for 15 minutes at the same flow rate. Residual NHS groups were quenched with ETA-HCl. IA@MoNPs, MoNPs, and bare NPs were injected sequentially over the VCAM1-coated surface, and SPR signals were recorded over a 400-s association phase and a 600-s dissociation phase. Kinetic parameters and affinity values were extracted using Scrubber 2.0c.

### 2.5. Nanoparticle uptake assays

ECs were pretreated with atheroprone stimuli, including TNFα (10 ng/mL, 3 hours), oxLDL (100 μg/mL, 24 hours), and low SS (1 dyne/cm^2^), prior to incubation with fluorescently labeled IA@MoNPs or MoNPs. For SS experiments, nanoparticle uptake was assessed using two approaches: post-SS, in which ECs were subjected to 24 hours of high or low SS before nanoparticle incubation, and under-SS, in which TNFα-primed ECs were exposed to nanoparticles under high or low SS. After incubation, unbound nanoparticles were removed, and cells were imaged and quantified for intracellular fluorescence using ImageJ. Mechanistic studies included blocking VCAM1 on TNFα -activated ECs with anti-VCAM1 (Cell Signaling #13622, 1:100), blocking β1-integrin on IA@MoNPs with anti-β1-integrin (Cell Signaling #4706S, 1:100), and inhibiting receptor-mediated endocytosis using chlorpromazine (CPZ) (Sigma #C8138-5G, 5 µM).

### 2.6. Animal experiments

All animal procedures were approved by the Institutional Animal Care and Use Committee (IACUC) of Arizona State University, USA. ApoE^□/ □^ mice, originally acquired from The Jackson Laboratory, were subjected to partial ligation (PL) of the left carotid artery (LCA) and then placed on a high-fat diet (HFD, Envigo #TD.88137) to induce endothelial activation and acute atherosclerosis [28, 16, 17]. Ten days post-PL, mice received 300 μg (dry weight) of fluorescently labeled IA@MoNPs or MoNPs via retro-orbital injection. Four hours later, arterial tissues, from the aortic arch to the carotid bifurcation, along with major organs, were harvested, fixed, and analyzed for fluorescent intensity using an *in vivo* imaging system (IVIS, PerkinElmer) at ASU Preclinical Imaging Core Facility. Following IVIS imaging, vessels were cryosectioned and subjected to fluorescent imaging to visualize the selective targeting of vascular lesions.

For circulation stability studies, C57BL/6 mice were intravenously injected with 300 μg of fluorescently labeled IA@MoNPs or MoNPs, and small volumes of blood were collected via the submandibular vein at 3- and 24-hours post-injection for IVIS imaging. Subsequently, whole blood obtained by cardiac puncture and major organs were harvested to assess acute toxicity. Blood chemistry was analyzed by Vetek Labs (Scottsdale, AZ), and complement activation was assessed by measuring serum C3a and C5a levels using ELISA kits (AFG Bioscience #CHM01J344 and #CHM01J346). Livers, kidneys, and spleens were cryosectioned and stained with hematoxylin and eosin (H&E) to evaluate histopathological changes.

### 2.7. Nanoparticle-blood interaction assays

IA@MoNPs or MoNPs were incubated with serum isolated from C57BL/6 mice at 37°C for 30 minutes to assess protein corona formation. Following incubation, nanoparticles were separated by centrifugation, and changes in their size were measured by DLS. The adsorbed protein corona was analyzed by Western blot using antibodies against complement 3 (C3) (Abcam #ab200999, 1:1000), immunoglobulin G (IgG) (Jackson ImmunoResearch #anti-rabbit IgG secondary antibody, 1:1000), ApoE (Cell Signaling #49285, 1:1000), and CD11b (Cell Signaling #17800, 1:1000). To evaluate nanoparticle clearance by phagocytes, mouse anti-coagulated whole blood was incubated with fluorescent IA@MoNPs or MoNPs at 37 °C for 30 minutes, followed by centrifugation to separate blood cells and plasma. Residual nanoparticle-associated fluorescence in the plasma fraction was quantified using IVIS imaging.

### 2.8. Statistical analysis

All data were derived from at least three independent experiments and are presented as mean ± standard deviation (SD). Student’s t-test and one-way analysis of variance (ANOVA) with Tukey’s multiple comparisons test were performed to evaluate statistical significance. Analyses were performed using GraphPad Prism software (GraphPad Prism 10), and a p-value < 0.05 was considered statistically significant.

## 3. Results

### 3.1. CCL2 or Mn^2+^ treatment induces VLA-4 activation and enhances IA@MoNP binding to VCAM1

To determine whether CCL2 or Mn^2^□ treatment induces membrane integrin activation that enhances interactions with endothelial ligands, we first examined the adhesion of treated monocytes to TNFα-activated ECs. Fluorescently labeled monocytes were stimulated with CCL2 or MnCl_2_, followed by incubation with inflamed EC monolayer. Microscopic analysis revealed that both treatments significantly increased monocyte adhesion compared with PBS controls (Fig. 1A). Adhesion strength was evaluated using a flow-induced detachment assay. After allowing monocytes to bind ECs under static conditions, unbound cells were removed and the same FOVs were imaged following sequential increases in laminar SS (1, 6, and 12 dyne/cm^2^). At 1 and 6 dyne/cm^2^, 65.6% and 57.6% of CCL2-treated monocytes and 65.3% and 56.0% of Mn^2+^-treated monocytes remained adherent versus 48.3% and 41.3% in PBS controls; at 12 dyne/cm^2^, 53.6% (CCL2) and 49.6% (Mn^2+^) of monocytes persisted compared with only 34.3% of control monocytes, indicating that CCL2 or Mn^2+^ treatment significantly increases monocyte adhesion strength (Fig. 1B). To confirm integrin activation, VLA-4 activity was assessed using LDV-FITC, a probe that selectively binds the high-affinity conformation of VLA-4. CCL2- or Mn^2+^-treated monocytes were incubated with LDV-FITC and analyzed by flow cytometry. Treated monocytes exhibited markedly elevated LDV-FITC fluorescence compared with PBS controls, confirming robust VLA-4 activation on monocyte membranes (Supplementary Fig. 1).

**Fig. 1.**
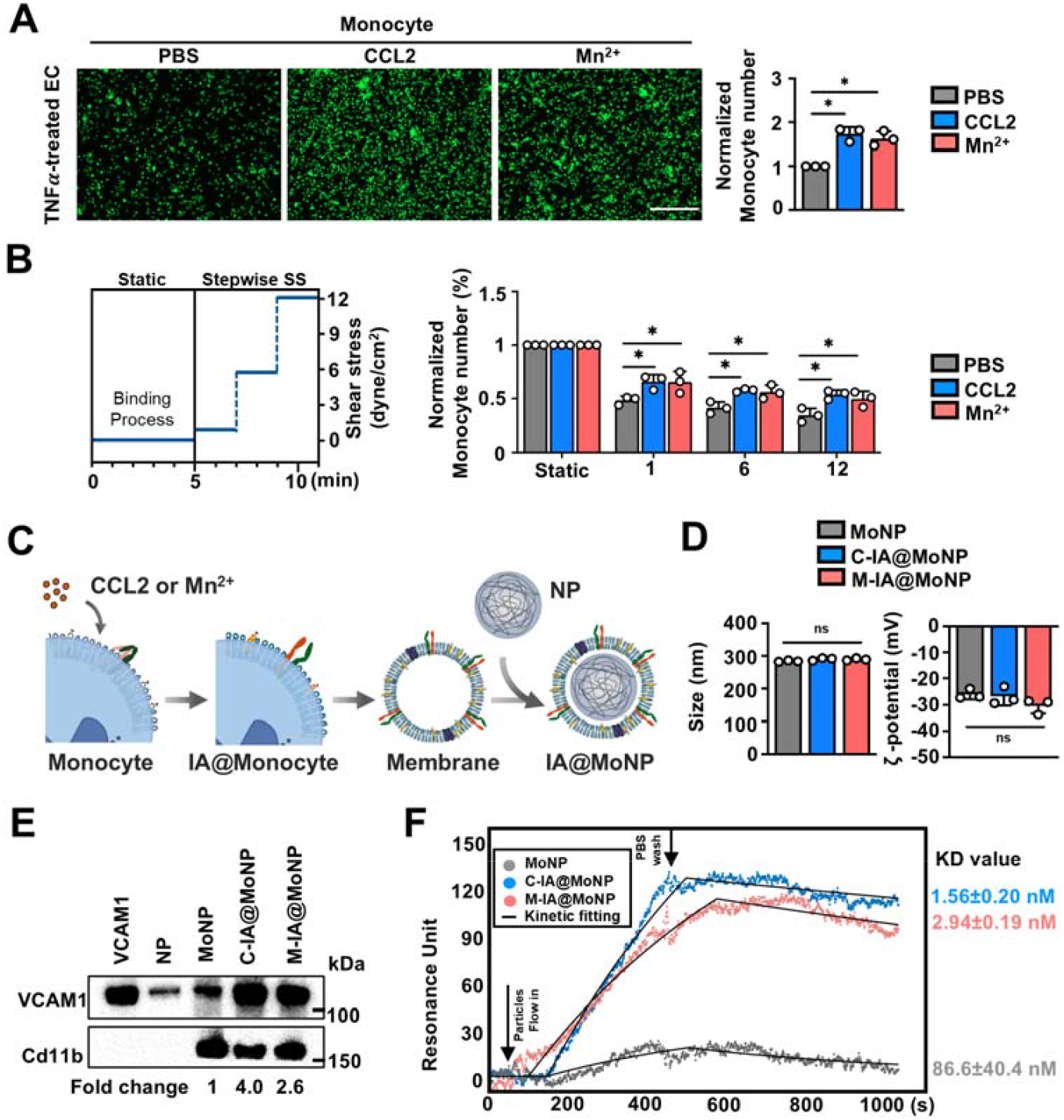
Membrane integrin activation enhances MoNP binding affinity to endothelial ligands. (A) Representative images and quantitative of adhesion of CCL2-, Mn^2+^-, or PBS-treated monocytes to TNFα-treated ECs. (B) Schematic of the monocyte adhesion assay under SS and quantification of fluorescently labeled monocyte attachment to TNFα-treated ECs. (C) Schematic illustration of C-IA@MoNPs and M-IA@MoNPs formulations. (D) DLS characterization showing hydrodynamic diameter and ζ-potential of IA@MoNPs. (E) Western blot and (F) SPR analyses demonstrating enhanced IA@MoNP binding affinity to recombinant VCAM1. CD11b was used as a loading control for all MoNP formulations. Data in (A) and (B) were normalized to PBS-treated monocytes and static conditions, respectively. (A–B) *p < 0.05 vs. PBS-treated monocytes. For all experiments, n = 3 independent replicates.

To assess whether activated integrins can be preserved and transferred onto nanoparticles, monocytes were stimulated with CCL2 or Mn^2+^ prior to plasma membrane isolation, and the resulting membranes were used to formulate C-IA@MoNPs and M-IA@MoNPs, respectively, following our established protocol (Fig. 1C) [16, 17]. DLS measurements showed that IA@MoNPs maintained comparable size and zeta potential to standard MoNPs prepared from naïve monocyte membranes (Fig. 1D). Western blot further confirmed the presence of CD11b and VLA-4 integrins (α4 and β1) on across all nanoparticle surfaces (Supplementary Fig. 2). We next evaluated whether integrin activation enhanced nanoparticle binding to the endothelial ligand VCAM1. IA@MoNPs or MoNPs were incubated with recombinant VCAM1 and analyzed by Western blot. Both IA@MoNP groups exhibited markedly stronger VCAM1 binding than MoNPs (Fig. 1E), indicating increased affinity driven by VLA-4 activation. Quantitative binding analysis using a modified SPR assay showed that IA@MoNPs produced substantially higher binding signals than MoNPs, with bare NPs exhibiting the lowest response (Fig. 1F and Supplementary Fig. 3). The equilibrium dissociation constants (K_D_) were 1.56 ± 0.20 nM for C-IA@MoNPs and 2.94 ± 0.19 nM for M-IA@MoNPs, compared with 86.6 ± 40.4 nM for MoNPs and >1 µM for bare NPs, confirming that IA@MoNPs exhibit roughly 30-50-fold higher binding affinity for VCAM1.

### 3.2. Membrane integrin activation enhances selective uptake by inflamed ECs

Having demonstrated that IA@MoNPs exhibit enhanced binding to VCAM1, we next evaluated their interaction with inflamed ECs. Three atheroprone stimuli—TNFα stimulation, oxLDL treatment, and prolonged low SS (1 dyne/cm^2^), a key hemodynamic component of disturbed flow—were examined, as each is known to induce EC activation and contribute to plaque development [29–32]. ECs were exposed to TNFα for 3 hours, oxLDL for 24 hours, or low SS for 24 hours; ECs subjected to high SS (12 dyne/cm^2^) served as the physiological shear control for the low SS condition. Western blot confirmed that all three atheroprone stimuli significantly upregulated VCAM1 expression compared with their respective controls (Fig. 2A), establishing an inflamed EC phenotype for IA@MoNP targeting studies. To assess nanoparticle uptake, DiD-labeled IA@MoNPs or MoNPs were incubated with inflamed ECs. Under TNFα stimulation, both IA@MoNP formulations showed significantly greater uptake than MoNPs, whereas no differences were observed among nanoparticle groups in control ECs (Fig. 2B). Similarly, IA@MoNPs exhibited enhanced binding to oxLDL-treated ECs compared with MoNPs, while all nanoparticle groups showed minimal uptake in control (Fig. 2C). We next evaluated nanoparticle adhesion under shear flow conditions. ECs were preconditioned under low SS or high SS for 24 hours to model atheroprone and atheroprotective conditions, respectively, prior to nanoparticle incubation. MoNPs exhibited greater adhesion to ECs following low SS exposure compared with high SS. Notably, IA@MoNPs further amplified adhesion to ECs under low SS relative to MoNPs, with no difference observed under high SS (Fig. 2D). To assess enhanced adhesion under active flow, nanoparticles were perfused over TNFα-activated ECs under low or high SS. IA@MoNP adhesion remained significantly higher than MoNPs under both shear conditions, with the strongest effect under low SS (Fig. 2E). Together, these findings demonstrate that integrin activation enhances MoNP selectivity for inflamed ECs without increasing nonspecific interactions with healthy ECs.

**Fig. 2.**
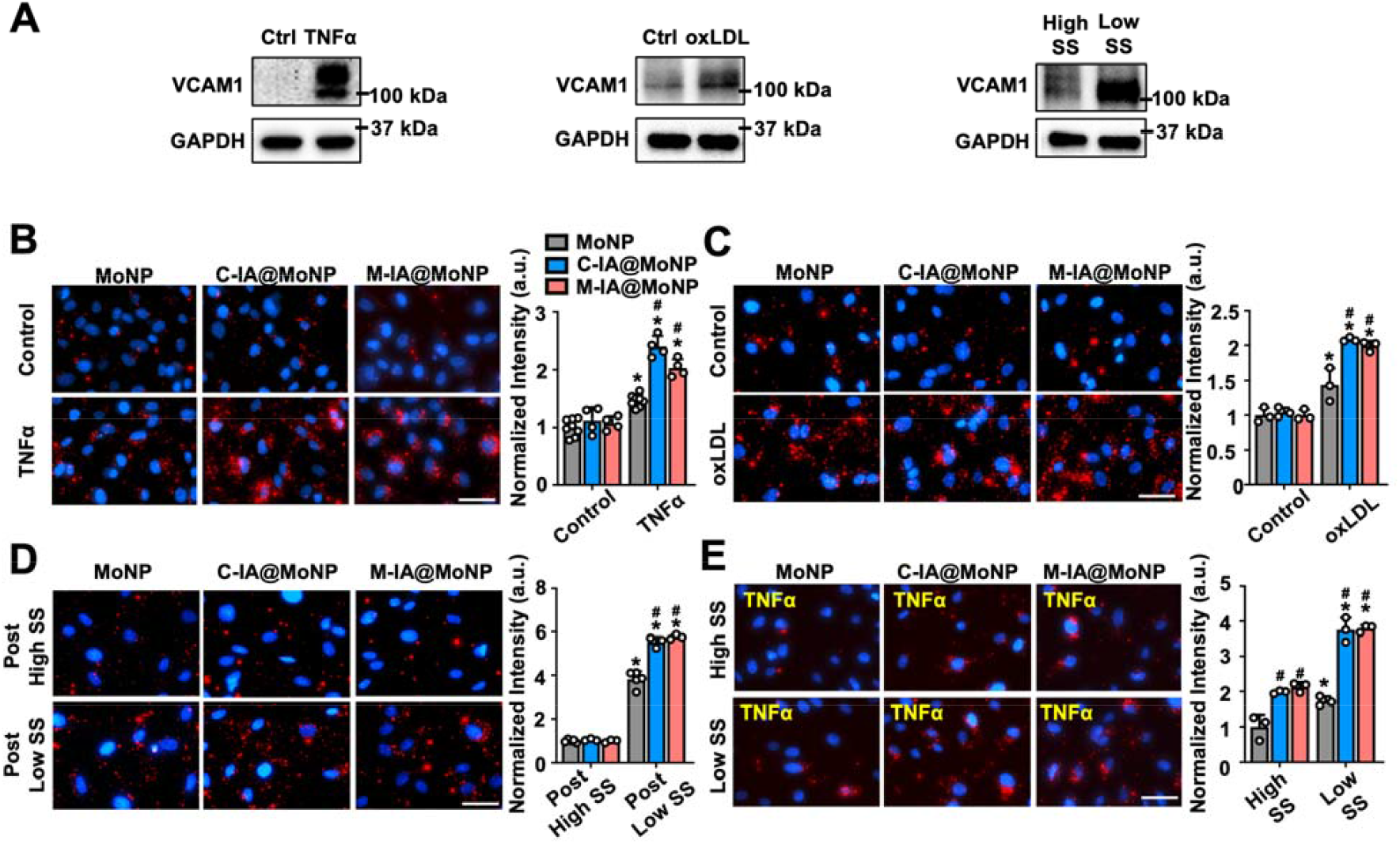
IA@MoNPs exhibit enhanced targeting of inflamed ECs *in vitro*. (A) Western blot analysis of VCAM1 expression in ECs exposed to TNFα, oxLDL, or 24-hour of shear flow. (B-E) Fluorescence images and quantification of nanoparticle uptake under the following conditions: (B) TNFα-activated ECs, (C) oxLDL-treated ECs, (D) ECs preconditioned with 24-hour shear flow followed by nanoparticle exposure, and (E) TNFα-activated ECs exposed to nanoparticles under continuous shear flow for 1 hour. Red: IA@MoNP or MoNP; blue: DAPI-stained nuclei. Scale bar = 50 μm. *p < 0.05 vs. control ECs; ^#^p < 0.05 vs. MoNP. Data are presented as mean ± SD from at least n = 3 independent experiments.

### 3.3. VLA-4/VCAM1 interaction governs IA@MoNP binding and uptake

We next investigated the molecular mechanisms underlying IA@MoNP interaction with inflamed ECs, focusing on the VLA-4/VCAM1 axis. To determine if binding was receptor-specific, we utilized a neutralizing antibody to block VCAM1 on the surface of inflamed ECs prior to incubation with C-IA@MoNPs, M-IA@MoNPs, or MoNPs. Fluorescence imaging and quantification showed that pre-treatment with anti-VCAM1 antibody—but not the IgG control—significantly reduced nanoparticle uptake across all formulations (Fig. 3A). To further confirm the requirement of the VLA-4/VCAM1 pair, we targeted the nanoparticle side of the interface by blocking β1-integrin, a key subunit of VLA-4 complex on the monocyte membranes. Consistently, β1-integrin blockade nearly abolished the endothelial uptake of both IA@MoNPs and MoNPs (Fig. 3B). Notably, the inhibitory effect of VCAM1 or β1-integrin blocking was markedly more pronounced for C-IA@MoNP and M-IA@MoNP groups compared to standard MoNPs. These results indicate that the VLA-4/VCAM1 pair is the primary driver of MoNP docking, with “activated” membranes providing a higher degree of the ligand-receptor engagement. To examine downstream internalization following surface docking, inflamed ECs were pretreated with CPZ, an inhibitor of clathrin-mediated endocytosis. CPZ significantly reduced intracellular nanoparticle accumulation (Fig. 3C), suggesting that VLA-4/VCAM1–mediated adhesion facilitates active, receptor-associated endocytic uptake rather than simple surface retention. Collectively, these data support a biomimetic mechanism in which activated monocyte membranes enhance endothelial targeting and promote subsequent nanoparticle internalization.

**Fig. 3.**
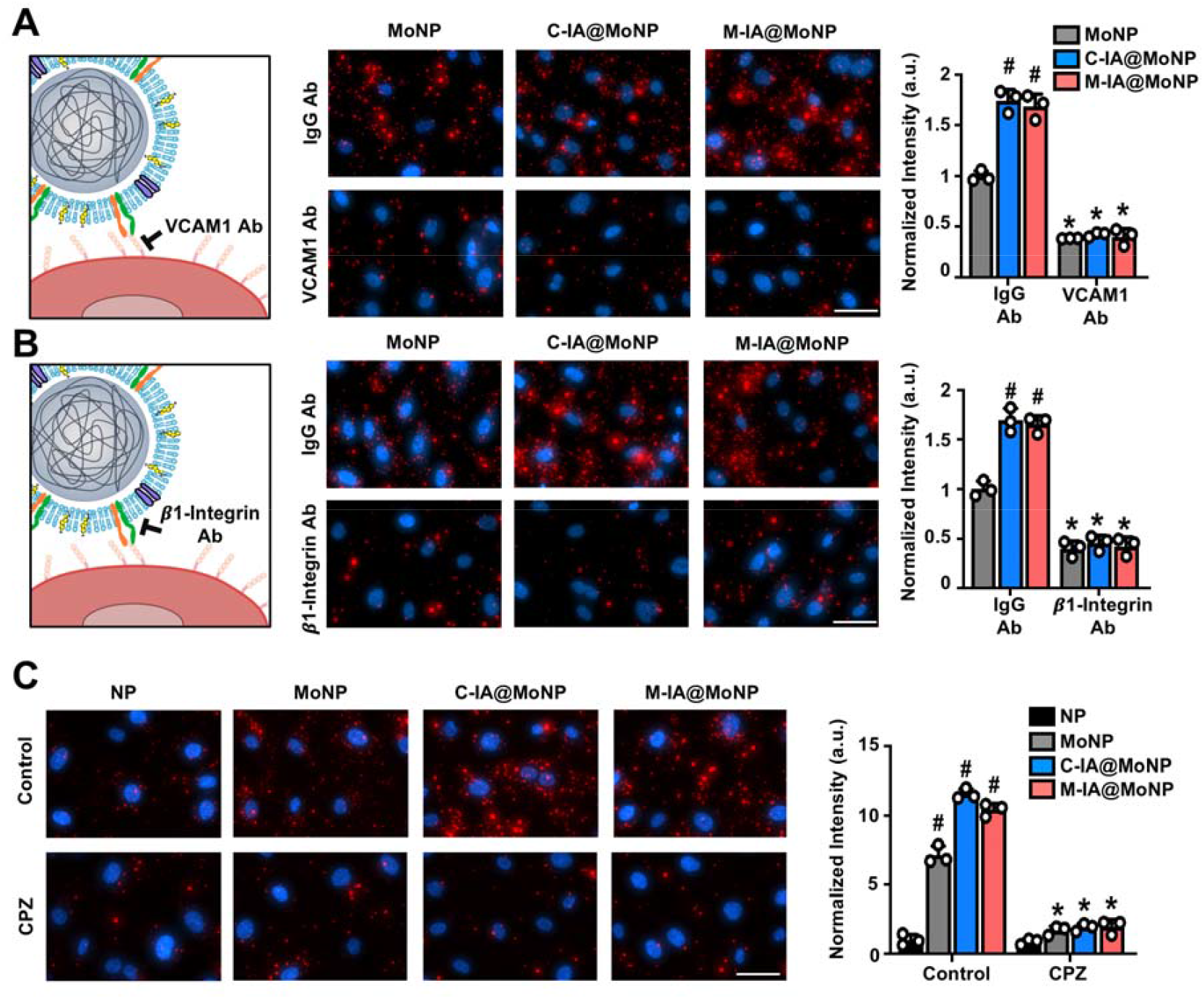
β1-integrin/VCAM1 axis mediates endocytic uptake of IA@MoNPs. Fluorescence images and quantification of nanoparticle uptake in TNFα-activated ECs under the following conditions: (A) VCAM1 blockade, (B) β1-integrin blockade on nanoparticles, and (C) CPZ treatment. Red: IA@MoNP, MoNP, or bare NP; blue: DAPI-stained nuclei. Scale bar = 50 μm. (A–B) *p < 0.05 vs. IgG antibody; ^#^p < 0.05 vs. MoNP. (C) *p < 0.05 vs. control ECs; ^#^p < 0.05 vs. bare NP. Data are presented as mean ± SD from n = 3 independent experiments.

### 3.4. IA@MoNPs exhibit enhanced targeting to atherosclerotic vessels *in vivo*

To evaluate the targeting performance of IA@MoNPs under complex physiological conditions, we employed a mouse model of disturbed flow-induced carotid atherosclerosis. ApoE^-/-^ mice were subjected to the PL procedure, followed by a HFD to induce rapid endothelial inflammation and plaque formation in the LCA [28, 16, 17]. Ten days post-procedure, when early-stage lesions were established, fluorescent C-IA@MoNPs, M-IA@MoNPs, or MoNPs were administered via retro-orbital injection. Four hours later, mice were euthanized, and vascular tissues and major organs were harvested for *ex vivo* IVIS imaging (Fig. 4A). The results revealed a markedly higher fluorescence signal in the atherosclerotic LCA of mice treated with IA@MoNPs compared to those treated with control MoNPs (Fig. 4B). Notably, this accumulation was highly specific to the inflamed LCA, with minimal fluorescence detected in the contralateral right carotid artery (RCA), aortic arch (AA), or descending aorta (DA). Cross-sectional tissue analysis further confirmed localized fluorescence within the LCA vessel wall for both IA@MoNP groups (Fig. 4C), indicating that integrin activation significantly enhances MoNP targeting to inflamed endothelium under disturbed flow. To confirm that targeting was driven by vascular inflammation, we performed parallel studies in control mice without the PL and HFD. No significant fluorescence was detected in the carotid arteries and aorta of these mice (Supplementary Fig. 4), indicating that IA@MoNP accumulation is driven by vascular inflammation. Furthermore, we assessed the systemic biodistribution of the formulations across major organs. IVIS analysis showed that the liver was the primary site of accumulation for all formulations, regardless of the membrane integrin activation (Fig. 4D). Importantly, membrane integrin activation did not increase non-specific binding in the heart, lungs, spleen, or kidneys compared to standard MoNPs. To test integrin-dependent targeting *in vivo*, we preincubated IA@MoNPs and MoNPs with a β1-integrin blocking antibody prior to administration. β1-integrin blockade markedly attenuated nanoparticle accumulation in the LCA (Fig. 4E and Supplementary Fig. 5). Collectively, these findings demonstrate that IA@MoNPs achieve superior and selective targeting of atherosclerotic arteries through a β1-integrin-dependent mechanism consistent with VLA-4-mediated adhesion, without increasing off-target accumulation to healthy tissues.

**Fig. 4.**
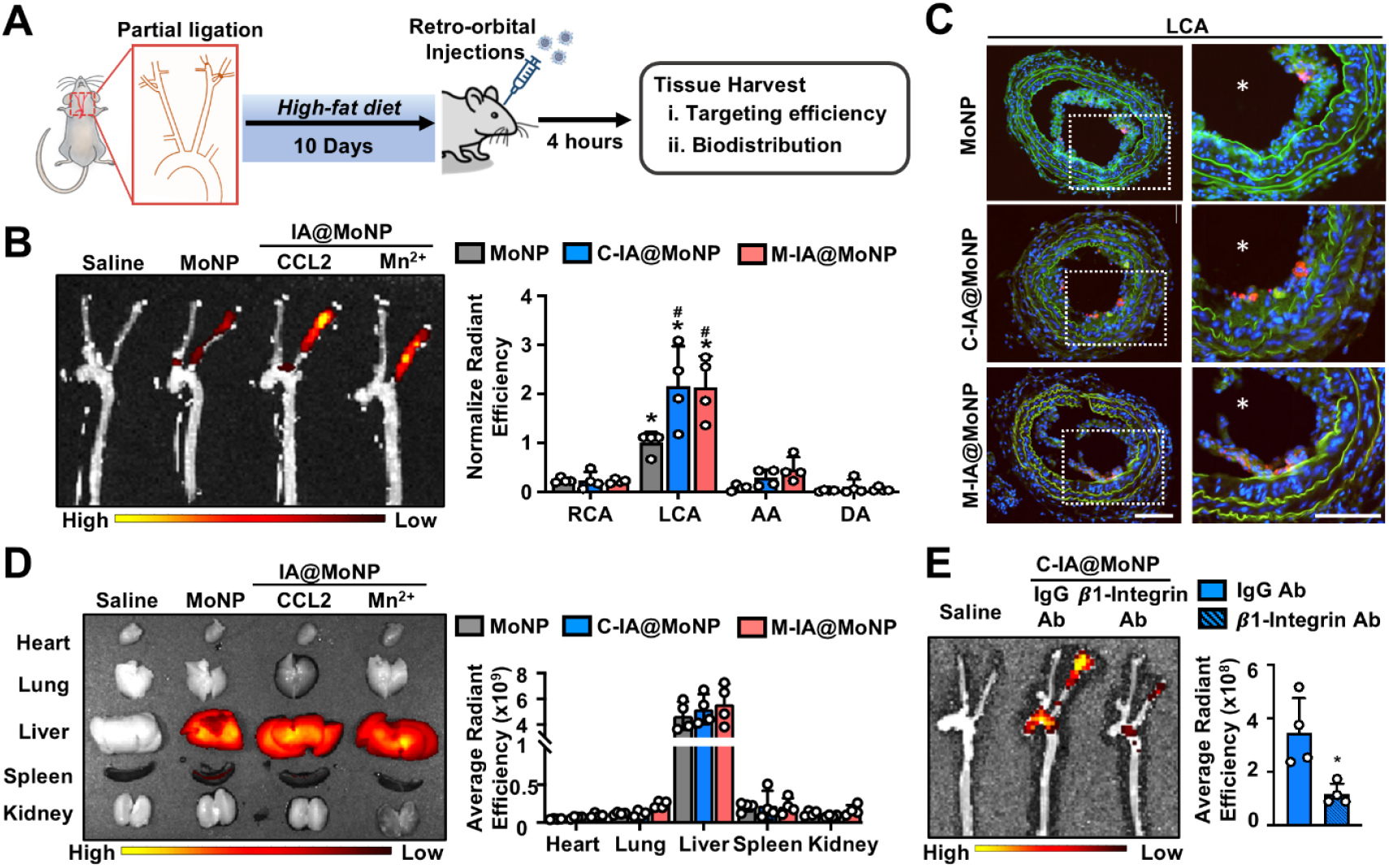
Integrin activation enhances MoNP targeting to atherosclerotic vessels. (A) Schematic illustration of the experimental design. (B) IVIS imaging and quantification of nanoparticle accumulation in partially ligated LCA of ApoE^-/-^ mice. (C) Representative cross-sectional fluorescence images of the LCA. Red: IA@MoNP or MoNP; green: elastin fibers; blue: DAPI-stained nuclei. Scale bar = 100 μm. (D–E) IVIS analysis showing (D) biodistribution in major organs across nanoparticle formulations and (E) reduction of IA@MoNP signal in the LCA following β1-integrin blockade on nanoparticles. (B) *p < 0.05 vs. RCA; ^#^p < 0.05 vs. MoNP; (E) *p < 0.05 vs. IgG antibody. Data are presented as mean ± SD from n = 4 mice each group.

### 3.5. Membrane integrin activation does not compromise biomimetic camouflage

Beyond enhanced inflammation targeting, a key advantage of monocyte membrane cloaking is the provision of biomimetic camouflage that improves circulation stability. A potential concern, however, is that conformational changes associated with integrin activation could alter the membrane physicochemical properties and compromise its “stealth” characteristics. To address this, we systematically evaluated nanoparticle stability, protein corona formation, and *in vivo* circulation profiles (Fig. 5A). We first assessed the colloidal stability of all nanoparticle formulations in serum. Following incubation with mouse serum for 30 minutes at 37°C, hydrodynamic size was measured via DLS. Bare NPs exhibited a clear increase in size, indicative of rapid protein adsorption and aggregation (Fig. 5B). In contrast, both IA@MoNPs and MoNPs maintained their original dimensions, demonstrating preserved colloidal stability in serum. To further characterize stealth performance, we analyzed protein corona composition, a critical determinant for mononuclear phagocyte system (MPS) clearance. Western blot analysis of nanoparticles retrieved from serum incubation showed that IA@MoNPs and MoNPs exhibited significantly lower levels of key opsonins, including C3, IgG, and ApoE [33], compared to bare NPs (Fig. 5C). These data suggest that the monocyte membrane cloaking serves as a barrier against common corona protein attachment, and importantly, integrin activation does not compromise the anti-opsonization function of the monocyte membrane coating. To evaluate blood compatibility, nanoparticles were incubated with anti-coagulated whole blood for 30 minutes at 37°C. IVIS imaging of the isolated plasma layer demonstrated significantly higher retention of both IA@MoNPs and MoNPs compared to bare NPs, with no discernible difference between the activated and standard membrane groups (Fig. 5D). Finally, *in vivo* circulation studies were performed in wild-type mice. Fluorescence measurements of blood samples collected at 3 and 24 hours post-injection revealed similar decay profiles for all membrane-cloaked formulations (Fig. 5E). No significant difference was observed between IA@MoNPs and MoNPs, indicating that integrin activation does not adversely affect systemic circulation time. Together, these findings highlight a key design advantage: membrane integrin activation enhances MoNP targeting without compromising the biological identity of the monocyte membrane coating.

**Fig. 5.**
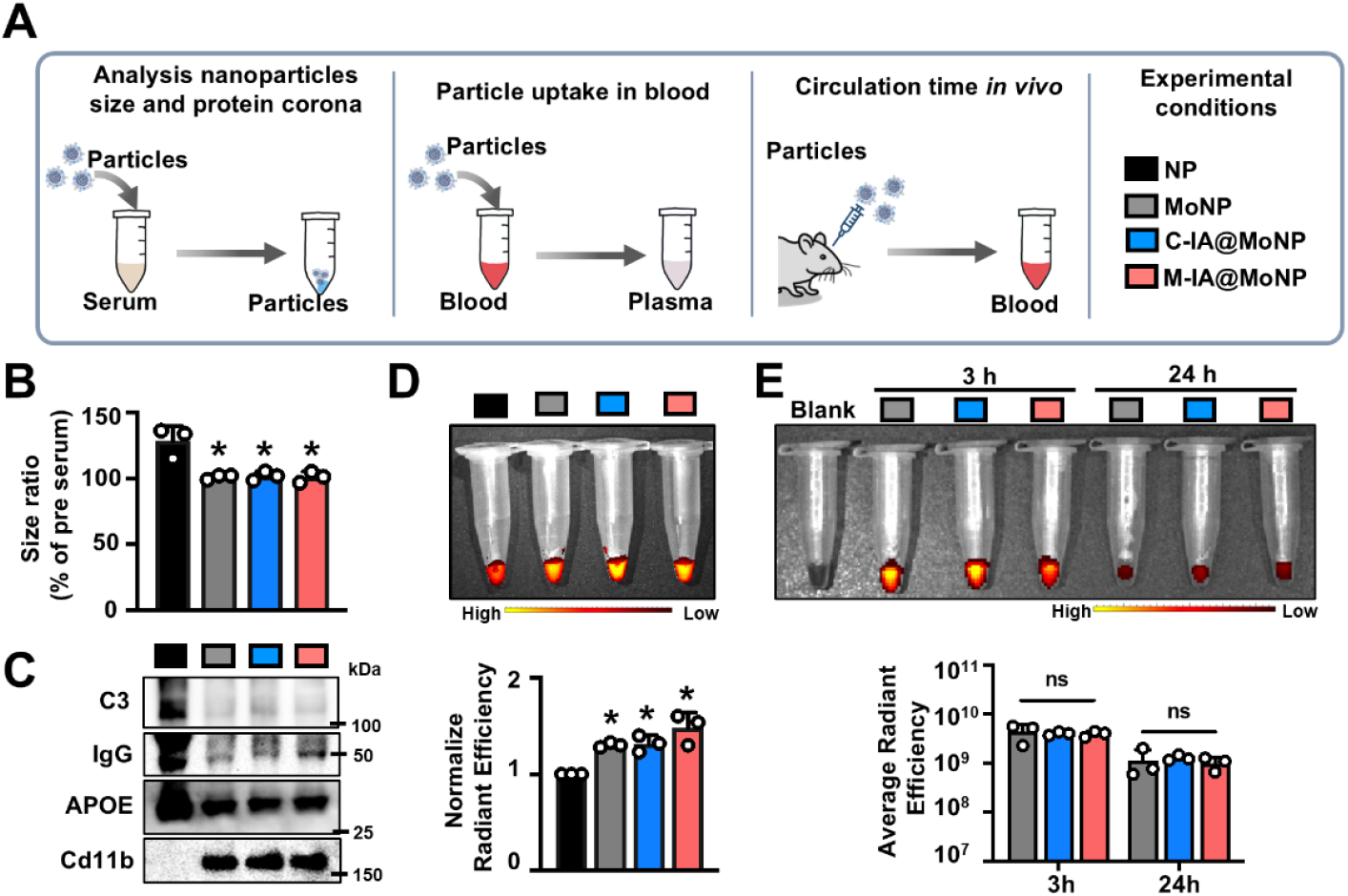
Membrane integrin activation does not alter the biomimetic properties of MoNPs. (A) Schematic illustration of the experimental design. (B–C) Nanoparticle characterization after serum incubation, showing (B) DLS analysis of hydrodynamic size and (C) protein corona composition by Western blot; CD11b was used as a loading control for all MoNP formulations. (D–E) IVIS images and quantification of residual nanoparticles, including (D) *ex vivo* measurement of nanoparticles remaining in plasma after incubation with whole blood and (E) *in vivo* measurement of circulating nanoparticles at 3- and 24-hour post-injection. (B, D): *p < 0.05 vs. bare NP. Data are presented as mean ± SD from n = 3 independent experiments.

### 3.6. Membrane integrin activation does not induce immunogenicity or systemic toxicity

Lastly, we evaluated the potential immunogenicity and systemic toxicity of the IA@MoNP formulations. Although the biosafety of MoNPs has been established in our previous studies [16, 17], it is essential to determine whether membrane integrin activation introduces new immunogenic epitopes or elicit acute inflammatory responses *in vivo*. A critical concern for intravenously administered nanoparticles is the potential for acute complement activation. To assess this, mice treated with IA@MoNPs or MoNPs were euthanized 24 hours post-injection. Serum samples were collected and analyzed for complement activation products C3a and C5a using ELISA. No significant differences in C3a or C5a were observed among IA@MoNP, MoNP, and saline groups (Fig. 6A). These results indicate that MoNPs do not trigger acute complement-mediated immune response, regardless of the membrane activation state. In parallel, blood samples were collected for comprehensive blood chemistry and metabolic analysis. As shown in Fig. 6B and Supplementary Fig. 6, no significant abnormalities were detected across a broad panel of metabolic markers. Specifically, biomarkers associated with hepatic function (e.g., ALT, AST) and renal function (e.g., creatinine, BUN) remained within normal physiological ranges across all treatment groups. These data indicate that neither the membrane cloaking nor the integrin activation process disrupts essential organ function. Moreover, histopathological examination of major metabolic organs, including liver, spleen, and kidneys revealed no evidence of inflammation or morphological abnormalities in any treatment group (Fig. 6C). Collectively, these findings demonstrate that IA@MoNPs are well-tolerated *in vivo*, supporting a robust safety profile for their applications in targeted drug delivery for atherosclerosis and other pathologies involving vascular inflammation.

**Fig. 6.**
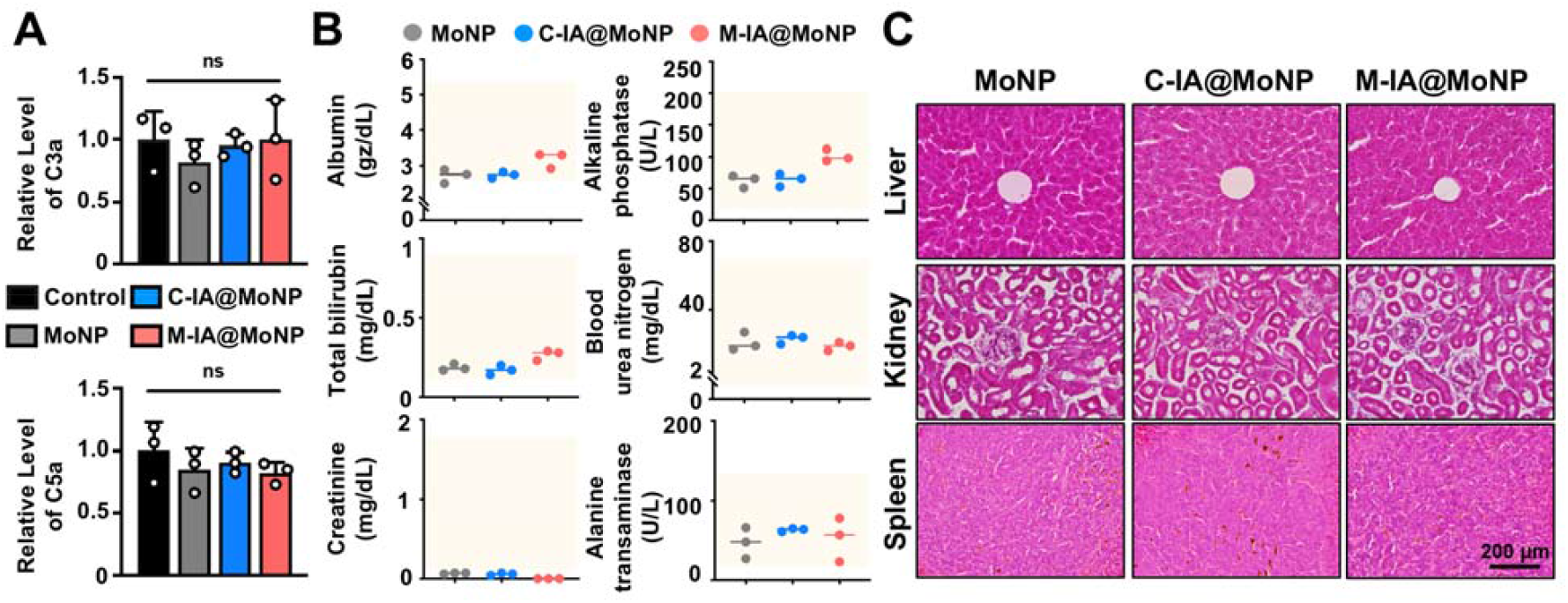
IA@MoNPs do not induce complement activation or organ toxicity. (A) Serum C3a and C5a levels measured by ELISA in mice injected with IA@MoNPs or MoNPs. (B) Serum metabolic panel analysis; yellow boxes indicate normal ranges. (C) Representative H&E staining of major organs from nanoparticle-treated mice. Scale bar = 200 μm. n = 3 mice per group.

## 4. Discussion

Ligand-conjugated nanoparticles that target endothelial adhesion molecules have emerged as a major strategy for vascular-targeted drug delivery, with peptides and antibodies against VCAM1 or ICAM1 among the most widely used approaches [34–37]. Leveraging cell membrane cloaking strategy, we established a MoNP platform that builds on this same biological principle but instead harnesses integrin receptors displayed on monocyte membranes to achieve site-specific delivery to atherosclerotic vessels. By preserving these integrins within their native lipid environment, monocyte membrane cloaking enables multivalent integrin receptor presentation, potentially enhancing binding avidity beyond what is achievable with single-ligand functionalization. This current study advances this platform by demonstrating that the targeting efficiency of MoNPs is governed not only by the presence of integrin receptors but, critically, by their affinity state. By engineering C-IA@MoNPs and M-IA@MoNPs with monocyte membranes pre-activated with CCL2 and Mn^2+^, respectively, we showed that the integrin receptors preserve their high-affinity conformational states, resulting in a 30-to-50-fold increase in binding affinity to VCAM1 (Fig. 1). Across multiple endothelial inflammation models, including TNFα stimulation, oxLDL exposure, and atheroprone flow, IA@MoNPs exhibit significantly enhanced uptake by inflamed ECs compared to standard MoNPs (Fig. 2). Mechanistically, this enhanced targeting is mediated by the VLA-4/VCAM1 interaction and subsequent receptor-mediated endocytosis (Fig. 1 and 3). Our *in vivo* findings in atherosclerotic mice confirmed that IA@MoNPs achieve significantly greater localization to lesions in the partially ligated carotid artery, without altering biodistribution across major organs (Fig. 4). Importantly, membrane integrin activation does not compromise the “stealth” properties, circulation stability, or safety profile of the original biomimetic platform (Fig. 5 and 6). These findings establish pre-activating membrane integrins as an effective approach to enhance the binding affinity and precision targeting of MoNPs to atherosclerotic vessels without compromising their biomimetic stealth and safety properties.

The critical role of VLA-4 integrin in vascular targeting has been demonstrated through engineered cell membrane systems [38–40]. For example, Park et al. showed that nanoparticles coated with membranes derived from genetically engineered murine leukemia cells expressing α4-integrin, which allows formation of the VLA-4 complex with endogenous β1-integrin, achieved significantly enhanced drug delivery to inflamed lung tissue compared those coated with unmodified membranes [39]. These findings established that membrane-displayed VLA-4 density contributed to VCAM1-mediated targeting to inflamed vasculature. While this genetic engineering approach increases VLA-4 abundance, our study introduces a complementary dimension: modulation of nanoparticle affinity by activating endogenous integrins prior to membrane isolation. To achieve integrin activation, we pretreated monocytes with either CCL2 or Mn^2+^, two mechanistically distinct approaches to inducing the high-affinity integrin conformational state. Both treatments produced IA@MoNPs with physicochemical properties comparable to standard MoNPs. However, our Western blot and SPR analyses confirmed that IA@MoNPs exhibit significantly enhanced binding to VCAM1, with 2.6-4-fold increase in protein association and a 30-50-fold enhancement in binding affinity. These results suggest that membrane integrin activation functionally primes the MoNP surface to recapitulate the high-affinity adhesive phenotype that monocytes adopt during the transition from rolling to firm endothelial adhesion, without altering nanoparticle physicochemical properties.

An essential requirement for vascular-targeted nanocarriers is the ability to distinguish diseased from healthy endothelium. Our results show that IA@MoNPs consistently exhibit enhanced targeting relative to standard MoNPs across multiple mechanistically distinct models of EC activation. This convergence suggests that membrane integrin activation enhances MoNP recognition of a shared adhesive EC phenotype characterized by elevated VCAM1 expression across diverse atheroprone stimuli. Importantly, increased IA@MoNP uptake remained restricted to inflamed ECs and was not observed in controls, indicating that integrin activation amplifies disease-selective targeting rather than promoting nonspecific endothelial adhesion. It is worth noting that the enhanced targeting of IA@MoNPs was preserved in ECs preconditioned under atheroprone flow but not under atheroprotective flow. Under active perfusion, IA@MoNPs exhibited greater binding than MoNPs, with particularly pronounced adhesion under low SS, suggesting integrin activation increases the overall binding strength of MoNPs while remaining effective particularly within disturbed flow regions where early lesions form. Together, these findings indicate that integrin activation improves MoNP sensitivity to the biochemical and hemodynamic cues in the atheroprone microenvironment, thereby enhancing targeting to atherosclerotic vasculature.

The enhanced targeting of IA@MoNPs was further supported *in vivo* using the PL mouse model, which generates a localized, highly atheroprone vascular environment characterized by low and oscillatory flow and consequent rapid plaque development [28]. Under these conditions, intravenously administered IA@MoNPs exhibited markedly greater accumulation in atherosclerotic regions compared with standard MoNPs, demonstrating that integrin activation enhances lesion-specific homing in complex physiological settings. Consistent with our *in vitro* findings, the increased binding affinity of IA@MoNPs remained highly selective in circulation and did not lead to elevated nonspecific vascular adhesion. Little to no signal was detected in non-inflamed vascular regions of PL mice or throughout the vasculature of control animals, indicating that enhanced targeting is dependent on the inflamed, adhesive endothelial phenotype. Notably, across both *in vitro* and *in vivo* models, IA@MoNPs consistently demonstrate substantially greater accumulation than MoNPs, supporting that integrin activation enhances targeting primarily by increasing receptor affinity rather than their abundance. Systemic biodistribution profiles were also preserved. The liver remained the primary site of nanoparticle accumulation, with no differences observed between IA@MoNPs and MoNPs. These findings indicate that the clearance pattern of IA@MoNPs remains consistent with standard MoNPs and the physiological trafficking and clearance behavior of circulating monocytes.

Mechanistically, our *in vitro* data demonstrate that blocking either β1-integrin on the IA@MoNP surface or VCAM1 on inflamed ECs markedly reduces nanoparticle uptake, indicating that enhanced adhesion requires VLA-4/VCAM1 engagement at the nanoparticle-endothelial interface. *In vivo*, blocking β1-integrin on either IA@MoNPs or MoNPs similarly abolished nanoparticle accumulation in the atherosclerotic carotid artery, providing further evidence that vascular homing depends on the ligand-receptor interactions under physiological conditions. While these findings strongly support a dominant role for the VLA-4/VCAM1 interaction, integrin activation induced by CCL2 or Mn^2+^ is not specific to VLA-4. Both treatments function as broad integrin activators, and adhesion of IA@MoNPs to inflamed endothelium is therefore likely multivalent. Consequently, we cannot exclude potential synergistic contributions from other common integrin complexes preserved on the membrane, including VLA-5 (α5β1), Mac-1 (αMβ2), and LFA-1 (αLβ2) [23, 41]. These integrin complexes may engage additional ligands, such as ICAM1 and fibronectin, to stabilize initial binding. Nevertheless, the VLA-4/VCAM1 axis appears to function as the primary molecular anchor for the MoNP platform. By inducing a high-affinity conformational state in VLA-4, IA@MoNPs achieve the necessary binding strength for firm adhesion and subsequent internalization into ECs. This high-affinity engagement not only ensures lesion-specific accumulation under physiological flow but also facilitates the effective cellular uptake required for the delivery of therapeutic payloads.

A major advantage of MoNPs over conventional ligand-functionalized delivery systems is their biomimetic camouflage, which enhances immune evasion and prolongs circulation stability. A potential concern with integrin activation, however, is that increasing a MoNP’s adhesive capacity could inadvertently enhance nonspecific serum protein binding, promote opsonization, and accelerate clearance by the MPS. Our findings demonstrate that integrin activation represents a highly discrete surface modification that does not alter nanoparticle stability in serum or the membrane’s intrinsic stealth properties in circulation. IA@MoNPs exhibited protein corona compositions comparable to those of standard MoNPs, with both showing significantly less opsonin adsorption relative to bare NPs. Consistently, *ex vivo* phagocytosis assays and *in vivo* circulation studies revealed no significant differences between IA@MoNPs and MoNPs. These results indicate that membrane cloaking provides a robust protective interface and that integrin activation enhances targeting without compromising the protective biomimetic interface. Nevertheless, protein corona composition can differ substantially between *in vitro* and *in vivo* environments [42]. Future studies using IA@MoNPs retrieved from serum incubation, followed by unbiased proteomic approaches, such as liquid chromatography-mass spectrometry, will provide deeper insight into corona composition and may guide further optimization for prolonged circulation and improved therapeutic efficacy.

Intravenously administered nanomedicine must avoid triggering immunogenicity and organ toxicity to enable clinical translation. Our comprehensive biosafety evaluation supports the translational feasibility of the IA@MoNP platform. The absence of elevated complement activation markers (C3a and C5a), together with normal hepatic and renal metabolic panels, indicates that IA@MoNPs, similar to standard MoNPs, are well tolerated and do not provoke systemic inflammatory responses. This safety profile is further supported by histological analysis of the liver, spleen, and kidneys, which revealed no evidence of morphological abnormalities. Despite these encouraging findings, several considerations remain for future development. First, although our data demonstrate a robust acute safety profile, additional studies are required to evaluate long-term safety following repeated administration. Second, the use of allogeneic membrane sources in preclinical models may not directly translate to human applications and remains a key challenge for clinical implementation. Future studies in larger animal models will be critical, as this risk may ultimately be mitigated by using autologous cell membranes derived from a patient’s own blood. Third, having established enhanced targeting efficiency, the next critical step is to evaluate optimal dosing strategies to minimize plaque progression in clinically relevant models and directly compare IA@MoNPs with standard MoNPs and systemic administration of free therapeutics. Finally, because EC activation and subsequent monocyte recruitment are central features of inflammatory vascular pathology across diverse organ systems, the high-affinity targeting mechanism achieved through integrin pre-activation on MoNPs may warrant evaluation beyond atherosclerosis.

## 5. Conclusion

In summary, we developed a simple and robust strategy to enhance vascular targeting efficiency by preactivating integrins on the surface of MoNPs. This modification increases binding affinity to the endothelial adhesion molecule VCAM1 on inflamed ECs, thereby improving nanoparticle localization to atherosclerotic sites. Our findings demonstrate that integrin activation significantly enhances adhesion to inflamed ECs through the VLA-4/VCAM1 axis, resulting in greater accumulation within atherosclerotic arteries compared with standard MoNPs. Notably, this strategy does not increase nonspecific interactions with healthy vasculature, compromise circulation stability, or induce organ toxicity. Collectively, these results establish integrin activation as a mechanistically driven approach to amplify MoNP targeting of vascular inflammation while preserving immune compatibility and systemic safety, supporting its potential theranostic applications in atherosclerosis and other inflammatory vascular diseases.

## Data availability statement

All date supporting the findings and/or analyzed during the current study are included within this article and its associated supplementary materials.

## Contributions

T.W. and K.W. conceived the study, performed data analysis, and wrote the manuscript. T.W. and J.R. conducted the experiments. K.H. assisted with methodology and investigation. J.J., Z.W., and S.W. contributed to the SPR experiments and data analysis. All authors reviewed and approved the final manuscript. K.W.

## Acknowledgements

The authors acknowledge the use of Regenerative Medicine Core and Preclinical Imaging Core facilities at Arizona State University.

## REFERENCES

1. Kong, P., Z.-Y. Cui, X.-F. Huang, D.-D. Zhang, R.-J. Guo, and M. Han. Inflammation and atherosclerosis: signaling pathways and therapeutic intervention. Sig Transduct Target Ther 7(1):131 2022. 10.1038/s41392-022-00955-7

2. Björkegren, J. L. M., and A. J. Lusis. Atherosclerosis: Recent developments. Cell 185(10):1630–1645 2022. 10.1016/j.cell.2022.04.004

3. Deanfield, J. E., J. P. Halcox, and T. J. Rabelink. Endothelial Function and Dysfunction: Testing and Clinical Relevance. Circulation 115(10):1285–1295 2007. 10.1161/CIRCULATIONAHA.106.652859

4. Gimbrone, M. A., and G. García-Cardeña. Endothelial Cell Dysfunction and the Pathobiology of Atherosclerosis. Circulation Research 118(4):620–636 2016. 10.1161/CIRCRESAHA.115.306301

5. Mitchell, M. J., M. M. Billingsley, R. M. Haley, M. E. Wechsler, N. A. Peppas, and R. Langer. Engineering precision nanoparticles for drug delivery. Nat Rev Drug Discov 20(2):101–124 2021. 10.1038/s41573-020-0090-8

6. Zhao, Z., A. Ukidve, J. Kim, and S. Mitragotri. Targeting Strategies for Tissue-Specific Drug Delivery. Cell 181(1):151–167 2020. 10.1016/j.cell.2020.02.001

7. Elumalai, K., S. Srinivasan, and A. Shanmugam. Review of the efficacy of nanoparticle-based drug delivery systems for cancer treatment. Biomedical Technology 5:109–122 2024. 10.1016/j.bmt.2023.09.001

8. Khodabandehlou, K., J. J. Masehi-Lano, C. Poon, J. Wang, and E. J. Chung. Targeting cell adhesion molecules with nanoparticles using in vivo and flow-based in vitro models of atherosclerosis. Exp Biol Med (Maywood) 242(8):799–812 2017. 10.1177/1535370217693116

9. Hsueh, P.-Y., Y. Ju, A. Vega, M. C. Edman, J. A. MacKay, and S. F. Hamm-Alvarez. A Multivalent ICAM-1 Binding Nanoparticle which Inhibits ICAM-1 and LFA-1 Interaction Represents a New Tool for the Investigation of Autoimmune-Mediated Dry Eye. Int J Mol Sci 21(8):2758 2020. 10.3390/ijms21082758

10. Dosta, P. et al. Delivery of Anti-microRNA-712 to Inflamed Endothelial Cells Using Poly(β-amino ester) Nanoparticles Conjugated with VCAM-1 Targeting Peptide. Adv Healthc Mater 10(15):e2001894 2021. 10.1002/adhm.202001894

11. Pickett, J. R., Y. Wu, L. F. Zacchi, and H. T. Ta. Targeting endothelial vascular cell adhesion molecule-1 in atherosclerosis: drug discovery and development of vascular cell adhesion molecule-1-directed novel therapeutics. Cardiovasc Res 119(13):2278–2293 2023. 10.1093/cvr/cvad130

12. Yu, M. et al. Nanoparticles targeting extra domain B of fibronectin-specific to the atherosclerotic lesion types III, IV, and V-enhance plaque detection and cargo delivery. Theranostics 8(21):6008–6024 2018. 10.7150/thno.24365

13. Muro, S. Challenges in design and characterization of ligand-targeted drug delivery systems. J Control Release 164(2):125–137 2012. 10.1016/j.jconrel.2012.05.052

14. Liu, H., Y.-Y. Su, X.-C. Jiang, and J.-Q. Gao. Cell membrane-coated nanoparticles: a novel multifunctional biomimetic drug delivery system. Drug Deliv. and Transl. Res. 13(3):716–737 2023. 10.1007/s13346-022-01252-0

15. Hu, C.-M. J. et al. Nanoparticle biointerfacing by platelet membrane cloaking. Nature 526(7571):118–121 2015. 10.1038/nature15373

16. Huang, H.-C. et al. Biomimetic nanodrug targets inflammation and suppresses YAP/TAZ to ameliorate atherosclerosis. Biomaterials 306:122505 2024. 10.1016/j.biomaterials.2024.122505

17. Rousseau, J. et al. Monocyte[Mimetic Contrast Agent Enables Targeted and Sensitive Magnetic Resonance Imaging of Atherosclerotic Lesions. Adv Healthcare Materials 15(3):e02001 2026. 10.1002/adhm.202502001

18. Liu, D.-Q. et al. Signal Regulatory Protein α Negatively Regulates β2 Integrin-Mediated Monocyte Adhesion, Transendothelial Migration and Phagocytosis. PLoS ONE 3(9):e3291 2008. 10.1371/journal.pone.0003291

19. Mestas, J., and K. Ley. Monocyte-Endothelial Cell Interactions in the Development of Atherosclerosis. Trends in Cardiovascular Medicine 18(6):228–232 2008. 10.1016/j.tcm.2008.11.004

20. Medrano-Bosch, M., B. Simón-Codina, W. Jiménez, E. R. Edelman, and P. Melgar-Lesmes. Monocyte-endothelial cell interactions in vascular and tissue remodeling. Front. Immunol. 14:1196033 2023. 10.3389/fimmu.2023.1196033

21. Ryu, J.-W. et al. Overexpression of Uncoupling Protein 2 in THP1 Monocytes Inhibits β <sub>2<sub> Integrin-Mediated Firm Adhesion and Transendothelial Migration. ATVB 24(5):864–870 2004. 10.1161/01.ATV.0000125705.28058.eb

22. Ashida, N., H. Arai, M. Yamasaki, and T. Kita. Distinct Signaling Pathways for MCP-1-dependent Integrin Activation and Chemotaxis. Journal of Biological Chemistry 276(19):16555–16560 2001. 10.1074/jbc.M009068200

23. Anderson, J. M., J. Li, and T. A. Springer. Regulation of integrin α5β1 conformational states and intrinsic affinities by metal ions and the ADMIDAS. MBoC 33(6):ar56 2022. 10.1091/mbc.E21-11-0536

24. Ye, F., C. Kim, and M. H. Ginsberg. Reconstruction of integrin activation. Blood 119(1):26–33 2012. 10.1182/blood-2011-04-292128

25. Davis, M. J., S. Earley, Y.-S. Li, and S. Chien. Vascular mechanotransduction. Physiological Reviews 103(2):1247–1421 2023. 10.1152/physrev.00053.2021

26. Koelbel, C., Y. Ruiz, Z. Wan, S. Wang, T. Ho, and D. Lake. Development of tandem antigen capture ELISAs measuring QSOX1 isoforms in plasma and serum. Free Radical Biology and Medicine 210:212–220 2024. 10.1016/j.freeradbiomed.2023.11.018

27. Yin, L. L., S. P. Wang, X. N. Shan, S. T. Zhang, and N. J. Tao. Quantification of protein interaction kinetics in a micro droplet. Review of Scientific Instruments 86(11):114101 2015. 10.1063/1.4934802

28. Nam, D. et al. Partial carotid ligation is a model of acutely induced disturbed flow, leading to rapid endothelial dysfunction and atherosclerosis. American Journal of Physiology-Heart and Circulatory Physiology 297(4):H1535–H1543 2009. 10.1152/ajpheart.00510.2009

29. Sawa, Y. et al. Effects of TNF-α on Leukocyte Adhesion Molecule Expressions in Cultured Human Lymphatic Endothelium. J Histochem Cytochem. 55(7):721–733 2007. 10.1369/jhc.6A7171.2007

30. Yurdagul, A., F. J. Sulzmaier, X. L. Chen, C. B. Pattillo, D. D. Schlaepfer, and A. W. Orr. Oxidized LDL induces FAK-dependent RSK signaling to drive NF-κB activation and VCAM-1 expression. Journal of Cell Science 129(8):1580–1591 2016. 10.1242/jcs.182097

31. Chiu, J.-J., and S. Chien. Effects of Disturbed Flow on Vascular Endothelium: Pathophysiological Basis and Clinical Perspectives. Physiological Reviews 91(1):327–387 2011. 10.1152/physrev.00047.2009

32. Walpola, P. L., A. I. Gotlieb, M. I. Cybulsky, and B. L. Langille. Expression of ICAM-1 and VCAM-1 and Monocyte Adherence in Arteries Exposed to Altered Shear Stress. ATVB 15(1):2–10 1995. 10.1161/01.ATV.15.1.2

33. Papini, E., R. Tavano, and F. Mancin. Opsonins and Dysopsonins of Nanoparticles: Facts, Concepts, and Methodological Guidelines. Front Immunol 11:567365 2020. 10.3389/fimmu.2020.567365

34. Muro, S., C. Gajewski, M. Koval, and V. R. Muzykantov. ICAM-1 recycling in endothelial cells: a novel pathway for sustained intracellular delivery and prolonged effects of drugs. Blood 105(2):650–658 2005. 10.1182/blood-2004-05-1714

35. Dosta, P. et al. Delivery of Anti-microRNA-712 to Inflamed Endothelial Cells Using Poly(β-amino ester) Nanoparticles Conjugated with VCAM-1 Targeting Peptide. Adv Healthc Mater 10(15):e2001894 2021. 10.1002/adhm.202001894

36. Zhou, Z. et al. Targeted polyelectrolyte complex micelles treat vascular complications in vivo. Proc. Natl. Acad. Sci. U.S.A. 118(50):e2114842118 2021. 10.1073/pnas.2114842118

37. Nahrendorf, M. et al. Noninvasive Vascular Cell Adhesion Molecule-1 Imaging Identifies Inflammatory Activation of Cells in Atherosclerosis. Circulation 114(14):1504–1511 2006. 10.1161/CIRCULATIONAHA.106.646380

38. Gorelik, M. et al. Use of MR Cell Tracking to Evaluate Targeting of Glial Precursor Cells to Inflammatory Tissue by Exploiting the Very Late Antigen-4 Docking Receptor. Radiology 265(1):175–185 2012. 10.1148/radiol.12112212

39. Park, J. H. et al. Genetically engineered cell membrane–coated nanoparticles for targeted delivery of dexamethasone to inflamed lungs. Sci. Adv. 7(25):eabf7820 2021. 10.1126/sciadv.abf7820

40. Jablonska, A. et al. Overexpression of VLA-4 in glial-restricted precursors enhances their endothelial docking and induces diapedesis in a mouse stroke model. J Cereb Blood Flow Metab 38(5):835–846 2018. 10.1177/0271678X17703888

41. Jiang, Y., J. F. Zhu, F. W. Luscinskas, and D. T. Graves. MCP-1-stimulated monocyte attachment to laminin is mediated by beta 2-integrins. American Journal of Physiology-Cell Physiology 267(4):C1112–C1118 1994. 10.1152/ajpcell.1994.267.4.C1112

42. Xiao, W. et al. The protein corona hampers the transcytosis of transferrin-modified nanoparticles through blood-brain barrier and attenuates their targeting ability to brain tumor. Biomaterials 274:120888 2021. 10.1016/j.biomaterials.2021.120888

