## Supplementary information for "Integrin Activation Enhances Lesion-Specific Targeting of Monocyte-Mimetic Nanoparticles in Atherosclerosis"

**Title**

501 E. Tyler Mall

Engineering Center G Wing, 346

Tempe, AZ 85287-9709

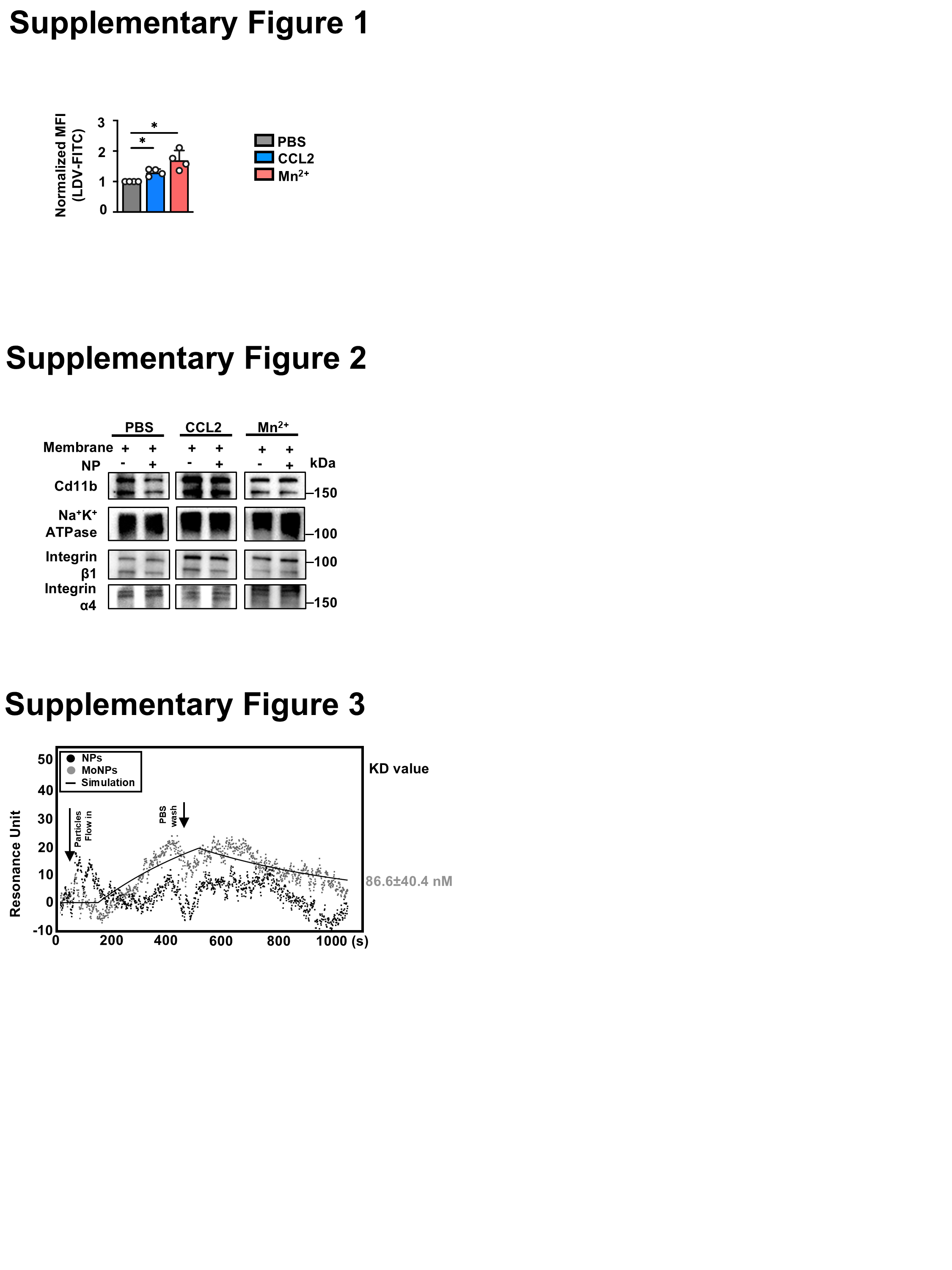

**Supplementary Fig. 1.** Mean fluorescence intensity (MFI) of LDV-FITC–labeled CCL2-, Mn²⁺-, or PBS-treated monocytes measured by flow cytometry. Data were normalized to the PBS group *p < 0.05 vs. PBS-treated group. Data are presented as mean ± SD from n = 4 independent experiments.

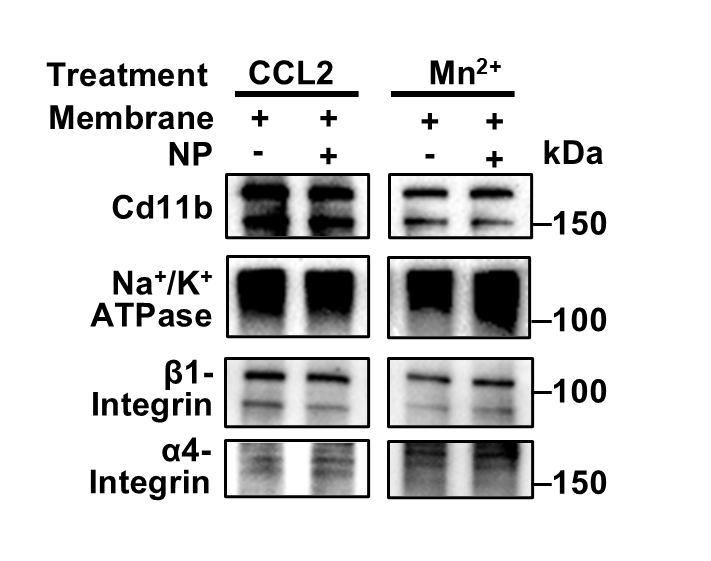

**Supplementary Fig. 2.** Western blot analysis of membrane proteins in IA@MoNPs.

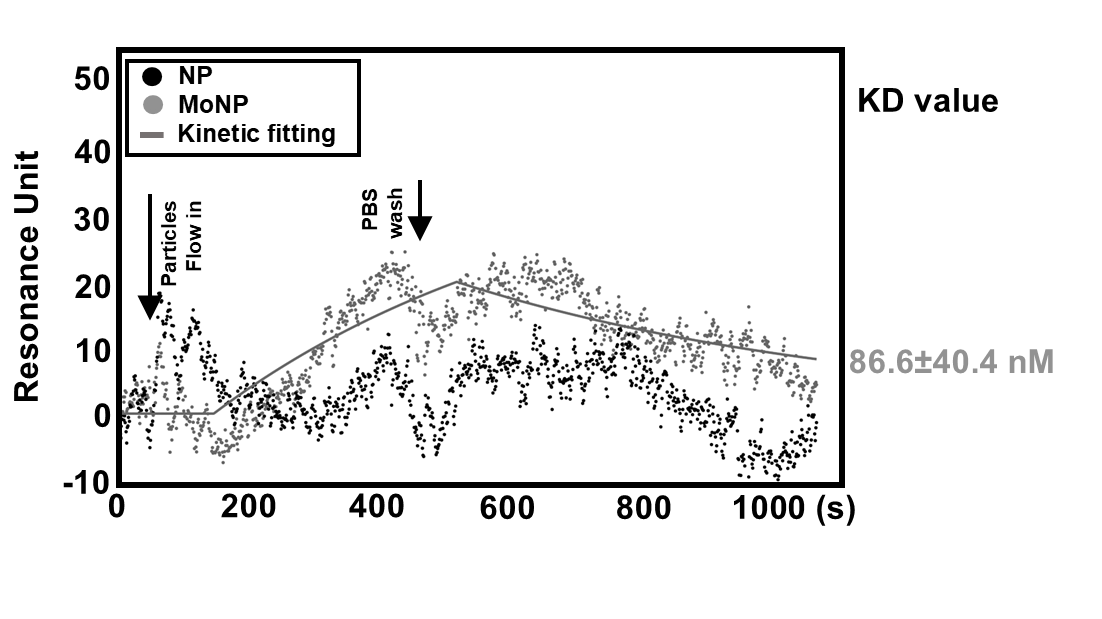

**Supplementary Fig. 3.** SPR analysis comparing the binding of MoNPs and bare NPs. n = 3 independent replicates.

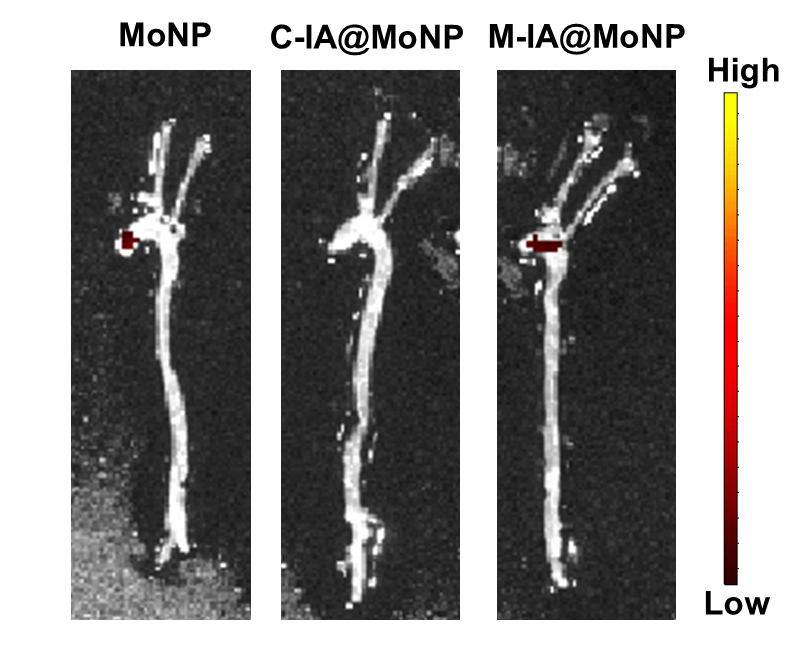

**Supplementary Fig. 4.** IVIS images of nanoparticle accumulation in control ApoE^-/-^ mice. n = 3 mice per group.

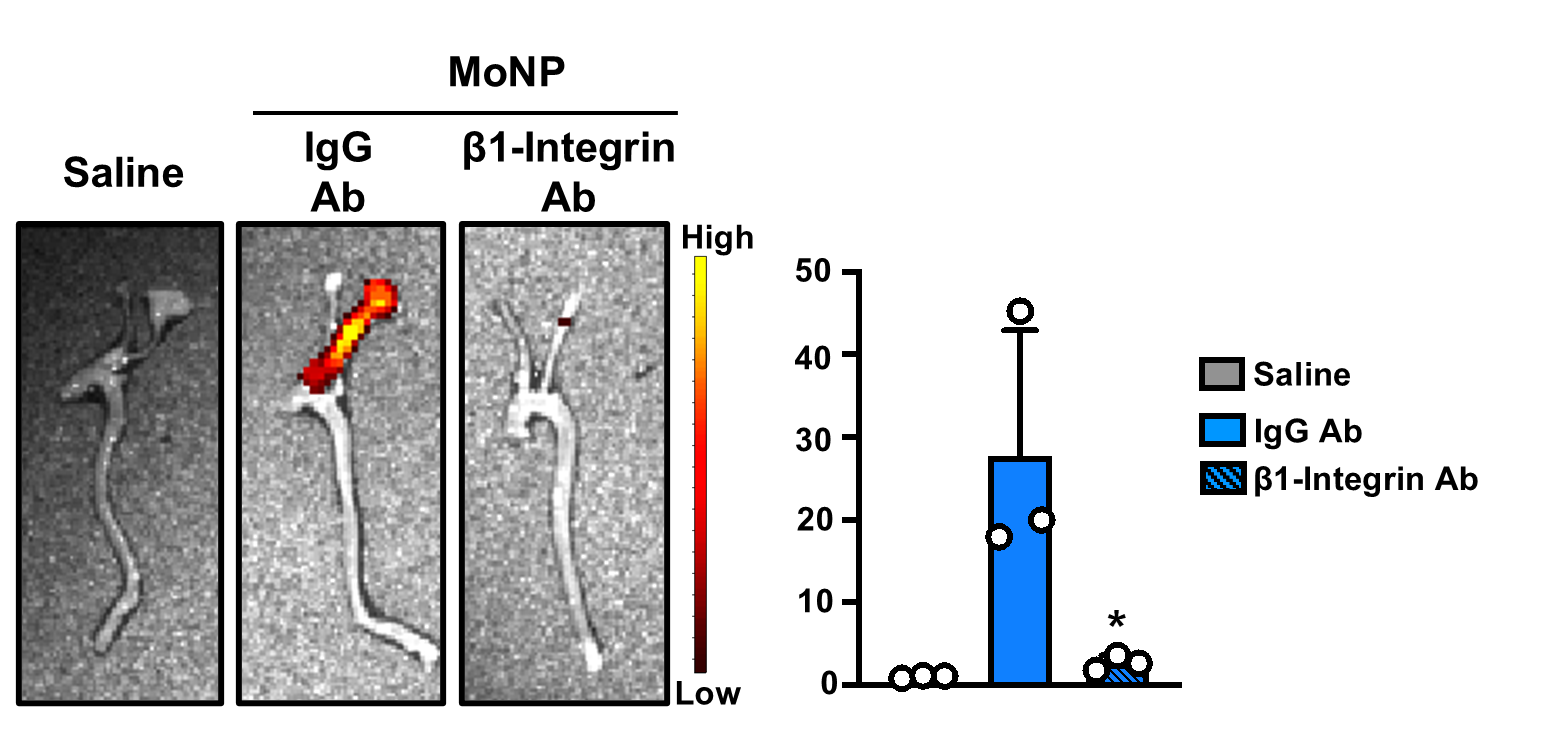

**Supplementary Fig. 5.** IVIS imaging and quantification of MoNP signal in the partially ligated left carotid artery following β1-integrin blockade on nanoparticles. *p < 0.05 vs. IgG. Data are presented as mean ± SD from n = 3 mice each group.

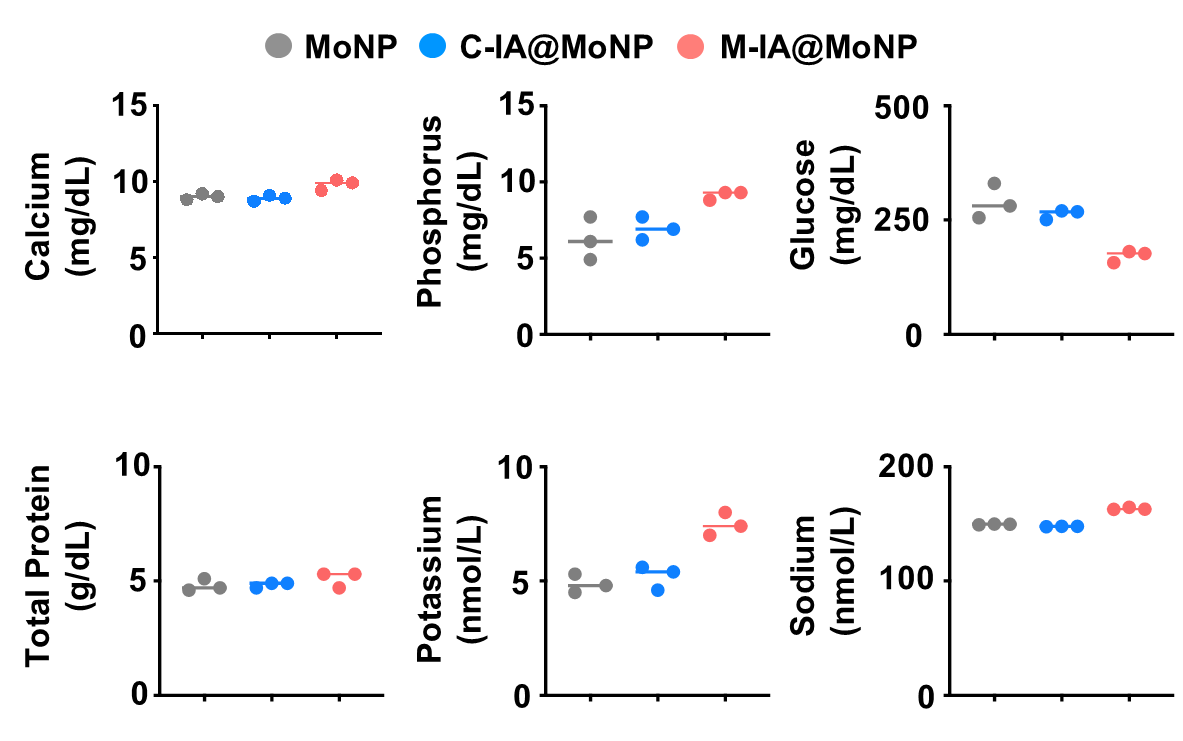

**Supplementary Fig.6.** Serum metabolic panel analysis of mice receiving IA@MoNPs and MoNPs. n = 3 mice per group.
